# Martini without the twist: Unveiling a mechanically correct microtubule through bottom-up coarse-graining in Martini 3

**DOI:** 10.1101/2024.05.29.596440

**Authors:** Abhilash Sahoo, Sonya M. Hanson

## Abstract

Microtubules are essential cytoskeletal filaments involved in cell motility, division, and intracellular transport. These biomolecular assemblies can exhibit complex structural behaviors influenced by various biophysical factors. However, simulating microtubule systems at the atomistic scale is challenging due to their large spatial scales. Here, we present an approach utilizing the Martini 3 Coarse-Grained (CG) model coupled with an appropriate elastic network to simulate microtubule-based systems accurately. By iteratively optimizing the elastic network parameters, we matched the structural fluctuations of CG hetero-dimer building blocks to their atomistic counterparts. Our efforts culminated in a ∼ 200nm microtubule built with ∼ 6 million interaction-centers that could reproduce experimentally observed mechanical properties. Our aim is to employ these CG simulations to investigate specific biophysical phenomena at a microscopic level. These microscopic perspectives can provide valuable insights into the underlying mechanisms and contribute to our knowledge of microtubule-associated processes in cellular biology. With MARTINI 3 CG simulations, we can bridge the gap between computational efficiency and molecular detail, enabling investigations into these biophysical processes over longer spatio-temporal scales with amino acid-level insights.

## Introduction

Microtubules are a dynamic and complex assembly of proteins, that form an essential component of the cellular cytoskeleton. These microscopic cylindrical structures along with other interaction partners contribute to an array of critical physiological functions within the cell such as maintaining cellular structure, supporting cell division, and facilitating intracellular transport. ^1,2^ Microtubules are an assembly of tubulin heterodimer subunits — *α*-*β* tubulin dimers that can rapidly assemble and disassemble, allowing cells to adapt to changing conditions and perform a wide array of dynamic functions.^3,4^ Structurally, these tubulins assemble into thin, hollow, cylindrical structures that polymerize from head-to-tail linearly creating protofilaments that in-turn can assemble into cylindrical assemblies with 11-16 protofilaments — with 13 being predominant in mammalian cells and 14 predominant *in vitro*. While *α* tubulins exclusively bind to GTP, *β* tubulins can exist in both GTP-bound and GDP-bound states *in vivo*. The conformation of tubulin dimers can vary based on their associated nucleotide states, leading to distinct structural features in the resulting microtubule assembly. Self-assembling microtubules can randomly switch between phases of growth and shrinkage, governed by these differences in structural properties — known as *dynamic instability*. Moreover individual tubulin dynamics along with their conformational variances, coupled with induced cooperativity due to the microtubule lattice structure, often result in highly dynamic assemblies that can respond to cellular and environmental stimulus allowing reorganization of cytoskeletal architecture in response. This presents a need for a bottom-up characterization of structure and dynamics of full microtubules starting from the microscopic details of individual dimer units.

Cryo-electron microscopy (cryo-EM) and X-ray crystallography have provided atomic resolution structures of various species of tubulin and microtubule structures with a variety of binding partners. ^5–8^ These binding partners have been shown to induce local structural diversity in tubulins and their assemblies. These features have been further correlated with macroscopic properties, dynamics and cellular function of microtubules. For example, recent experiments have reported how environmental stimulus such as buffer conditions and macromolecular crowding can influence microtubule biophysics such as mechanical properties, crowder-concentration induced bundling and interaction between microtubules. ^7,9–12^ To supplement the molecular resolution detail of structure determination and the experiments that give access to macroscopic biophysical properties, but can lack mechanistic insight, computational and theoretical tools have been employed to develop a fuller picture of microtubule biophysics. Examples of simple and intuitive *in silico* models of microtubules range from mean field approaches that do not explicitly incorporate detailed structural data to rule-based stochastic methods built with structural constraints from experimentally resolved structures. ^13–15^

Another computational approach that can be applied to understanding microtubules is molecular dynamics (MD) simulations. While MD can be a powerful computational technique to model complex structural assemblies while also accounting for atomic-level insights, the length scales required to model a full microtubule can make the method computationally prohibitive for this system. However, the structural characterization of tubulin dimers and smaller microtubule patches has been extensively studied with molecular simulations. ^16–20^ Applications of all-atom scale MD (AA-MD) is limited here to studies of dimers or dimer assemblies, less than 30-50 nm long. To our knowledge the largest relevant AA-MD study consists of a 14 protofilament microtubule with 6 dimers per protofilament, aiming to compare the dynamics of microtubule tips at different nucleotide associated states. ^19^ An appealing alternative approach however is coarse-grained molecular dynamics (CG-MD) simulations, developed to bypass the computational bottleneck of AA-MD by reducing the resolution and locally averaging structural and dynamical information. ^21^ This increases diffusion by decreasing local complexities in the free energy landscape and therefore allows extensive computational speed-up.

Modelling efforts along this vein aim to build larger microtubule structures from atomic level information, following a bottom-up coarse-graining approach, and allow bridging scales closer to experimental levels. For example, Kononova *et al.* built a bottom-up C*α*-based self organizing polymer (SOP) model with parameters derived from implicit solvent molecular dynamics with CHARMM19 force-field.^22^ This work established an *in-silico* nano-indentation method with results comparable to experimental force–deformation spectra and also revealed free energies of tubulin-tubulin interactions — both lateral and longitudinal. Another approach by Deriu *et al.* involved using molecular dynamics to refine protein conformations and normal mode analysis to extract vibrational characteristics, which was subsequently used to create a coarse-grained model. ^23^ However, these methods are non-transferable and rely on explicit parameterization of the whole system, limiting their usage in other physiological contexts beyond the direct setup or the problem it was parameterized for.

To address this gap, in this work we propose a coarse-grained biomolecular model for micro-tubules built with a commonly used and general coarse-grained forcefield — Martini 3. ^24^ Martini is an intermediate-level coarse-grained model with 1-5 interaction sites representing each amino-acid. This approach reduces computational demands while retaining essential structural and dynamic information. Martini has gained popularity for simulating large and complex biological systems, such as virions and organelles. Here, we report specific modifications to the elastic network architecture with the Martini 3 forcefield through an iterative bottom-up approach. Our refined intermediate-resolution coarse-grained model of the microtubule demonstrates the capability to map structural and dynamical data from accurate all-atom simulations to larger scale structures, while preserving the transferability of the forcefield. In this work, we have tested this approach on GDP-associated microtubule lattice, which due to low structural stability is difficult to isolate and study by experimental structural biology methods, and discussed the initial structural insights from this strategy.

## Results

### An initial coarse-grained microtubule proves unstable

In the following sections we discuss the modifications we employed to the Martini 3 forcefield towards creating a stable coarse-grained simulation of this microtubule.

### Building a complete microtubule lattice

Since microtubule structure and dynamics involves a complex interplay of tubulin dimers in various associated-nucleotide states, it is important to carefully prepare the initial lattice architecture. To build our Martini microtubule, we built our initial configuration according to the cryo-EM structure of undecorated microtubule with GDP-tubulin lattice (6DPV). ^25^ We extrapolated the microtubule’s inherent helical twist (Rotation -*PF_n_*/*PF_n_*_+1_, 0.874 nm) and axial rise (Z-rise -*PF_n_*/*PF_n_*_+1_, 25.77 degrees) from these segments to synthesize a single turn of the 14 protofilament-microtubule (Fig. 1A). Thereafter, these individual helical fragments were translated along the microtubule’s axis to create the initial microtubule structure with ∼ 25 dimers per protofilament. Note that the microtubule seam in this context was deduced through rotations and translations of protofilaments, so any potential distinct conformational variation of the tubulins at the seam is not present. We used the *martinize* protocol to map the all-atom microtubule structures to the coarse-grained Martini 3 representation. ^26^

**Figure 1:**
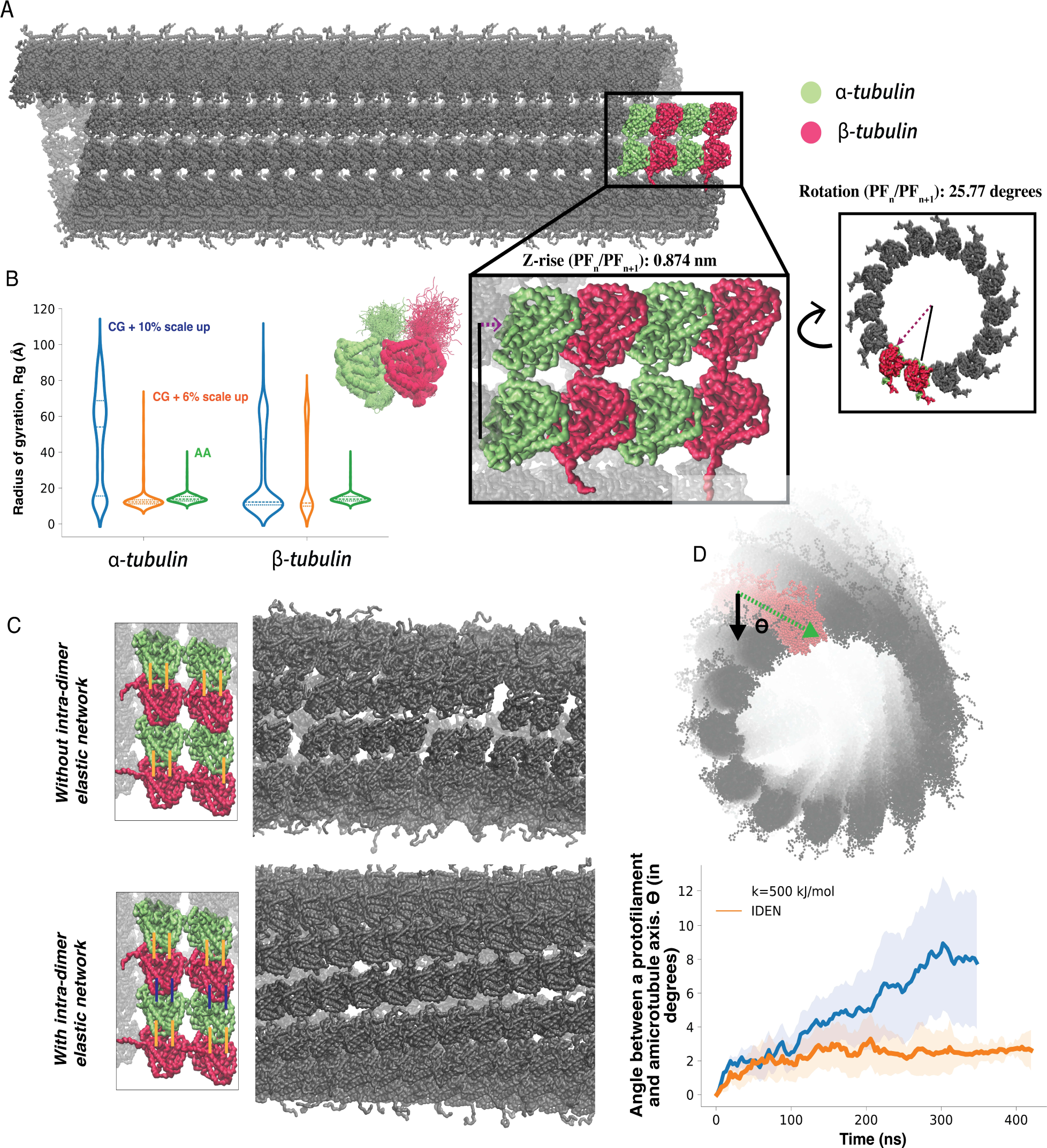
An initial CG microtubule. *(A)* Model CG microtubule (14-3) assembled from translat-ing and rotating cryo-EM structure (6DPV). *(B)* Radius of gyration (*R_g_*) of tubulin tails, compared across scaled protein-water interactions. *C* Out-of-box Martini 3 simulation with elastic net-work across a single dimer only (*top*), across the entire protofilament (*bottom*). *D, E* Twist in the protofilament due to constant elastic-network across the whole protofilament.

### Disordered C-terminal tails

One advantage of using the Martini coarse-grained model is that residue-level detail is preserved while achieving a computational speed up. For example, both *α*- and *β*-tubulins feature disordered and highly charged C-terminal tails 19 and 18 amino-acid long respectively, that can modulate their interactions with ions, metabolites, membranes, cytoskeletal assemblies, and microtubule-associated proteins (MAPs). ^27–30^ Previous studies have highlighted several alterations in the biophysical features and function of microtubules in the presence of intact C-terminal tails compared to those with truncated tails. ^31,32^ Therefore, it is necessary to model the C-terminal tails accurately, to allow this model to be extended to studies with a broader physiological focus. However, a prevalent challenge associated with Martini, is its tendency to promote more compact structural conformations, particularly in disordered regions. To mitigate this issue, a commonly employed strategy in both coarse-grained and atomistic forcefield development involves amplifying the interaction strength between protein and water.

Here, we adhered to the approach recommended by Thomasen *et al.*, wherein the authors proposed an *ad hoc* range of scaling factors for protein-water interactions from 6% to 10 % to effectively reproduce small-angle X-ray scattering (SAXS) data for multi-domain proteins. ^33^ This scaling factor was adopted to counterbalance the compacting effects of the forcefield and improve the accuracy of our simulations. To validate these modifications, we simulated a single GDP tubulin at atomistic resolution and compared against the mapped coarse-grained simulations, with up-scaled interaction by 6% and 10 %. We used radius of gyration (*R_g_*), which is a common validation metric for intrinsically disordered segments and was recently employed to parameterize a machine learning based forcefield for intrinsically disordered protein segments, to compare the structural ensemble (Fig. 1 B, C). ^34^ The distribution of *R_g_* for the reference atomistic simulation was unimodal, with a long tail and a median of 14.2Å. The 10% up-scaling from the Martini 3 forcefield resulted in a bi-modal distribution for both the tubulin types. But with 6% up-scaling we observed unimodal distribution for *α*-tubulin and bimodal distribution for *β*-tubulin, with the dominant population (median 13Å) overlapping with the atomistic simulation. Therefore, for the non-bonded interactions we have used the Martini 3 forcefield with the protein-water interactions up-scaled by 6%.

### Out-of-the-box Martini results in broken/twisted microtubule lattice

Since the Martini forcefield is limited in its ability to accurately capture the details of highly charged groups and small molecules such as the nucleotides like GTP and GDP, we decided not to explicitly include these nucleotides, but rather encode the structural and dynamical aspects of the nucleotide presence through a carefully designed elastic network model (ENM). Since the tubulin dimers constitute the fundamental units of microtubules, we initially applied an elastic network with a constant strength on the tubulin-dimers only. This preserves the initial GDP-tubulin structure, and relies on the protein-protein interactions from the Martini 3 forcefield to stabilize the overall lattice assembly. This ENM was designed with the default settings of *martinize*, with interaction sites defined as *in-contact*, if the distance between them ranged from 5Å to 9Å; with a spring constant of 500 kJ/mol. Also, as noted, since the C-terminal tails are known to be disordered, we excluded the them from the elastic-network. We ran 400 ns of this initial out-of-the-box CG microtubule and found that the microtubule lattice fell apart rapidly, resulting in a loss of most intra-dimer contacts by the last 200 ns (Fig. 1C *top*). This is probably due to the unstable protein-protein contacts at the inter-dimer interface resulting from the absence of explicit nucleotides to stabilize them.

In the next iteration we extended the ENM to include inter-dimer contacts, along a single protofilament, to maintain inter-dimer contacts and the fidelity of the microtubule lattice structure. With the new extended ENM, the dimer-dimer interactions were preserved by the network, preventing breaking up of the lattice (Fig. 1C *bottom*, Fig. 1D). However, the overall microtubule architecture started continuously twisting to maximize inter-protofilament contacts, nicely quantified by the angle that a protofilament makes with the microtubule axis (Fig. 1E).

While, this twisting could in theory be solved by extending the ENM to inter-protofilament contacts, we have avoided this approach in our systems as inter-protofilament contacts are much weaker than intra-protofilament contacts; and are not explicitly modulated by interactions with nucleotides/ligands. Moreover, not encoding the inter-protofilament contacts through an elastic network also allows extending these systems to studies involving microtubule tips, mechanical deformations and lattice defects.

To remove systematic effects that might arise due to the imbalance in inter-protofilament interactions at the tips, particularly near the microtubule seam, and create a more biologically relevant microtubule patch, we randomized the length of the protofilament by removing tubulin dimers at both the ends.

### Systematic optimization of the ENM

Here, we hypothesize that scaling down the elastic network strength would allow more micro-scopic local fluctuations and reduce the systematic twisting of the microtubule. But instead of an explicit scaling, with a refined elastic network, we can appropriately encode these local fluctuations from an atomistic representation onto our coarse-grained forcefield in a bottom-up manner. Our approach, with some careful alterations, closely follows the approach outlined by Globisch *et al.*, for capturing accurate structure and dynamics of the Cowpea Chlorotic Mottle Virus (CCMV) ^35^(Fig. 2B). First, we created an all-atom microtubule patch of three protofilaments, with four dimers each, built as previously described. The tubulins at the ends of the protofil-ament were center-of mass position restrained (10 kJ/mol) to simulate conditions within the microtubule lattice (Fig. 2A). The new ENM thus has variable network strength based on the all-atom simulations, characterized by a set of spring constants (K = {*k_ij_*}) and the corresponding equilibrium lengths (D = {*d_ij_*}).

**Figure 2:**
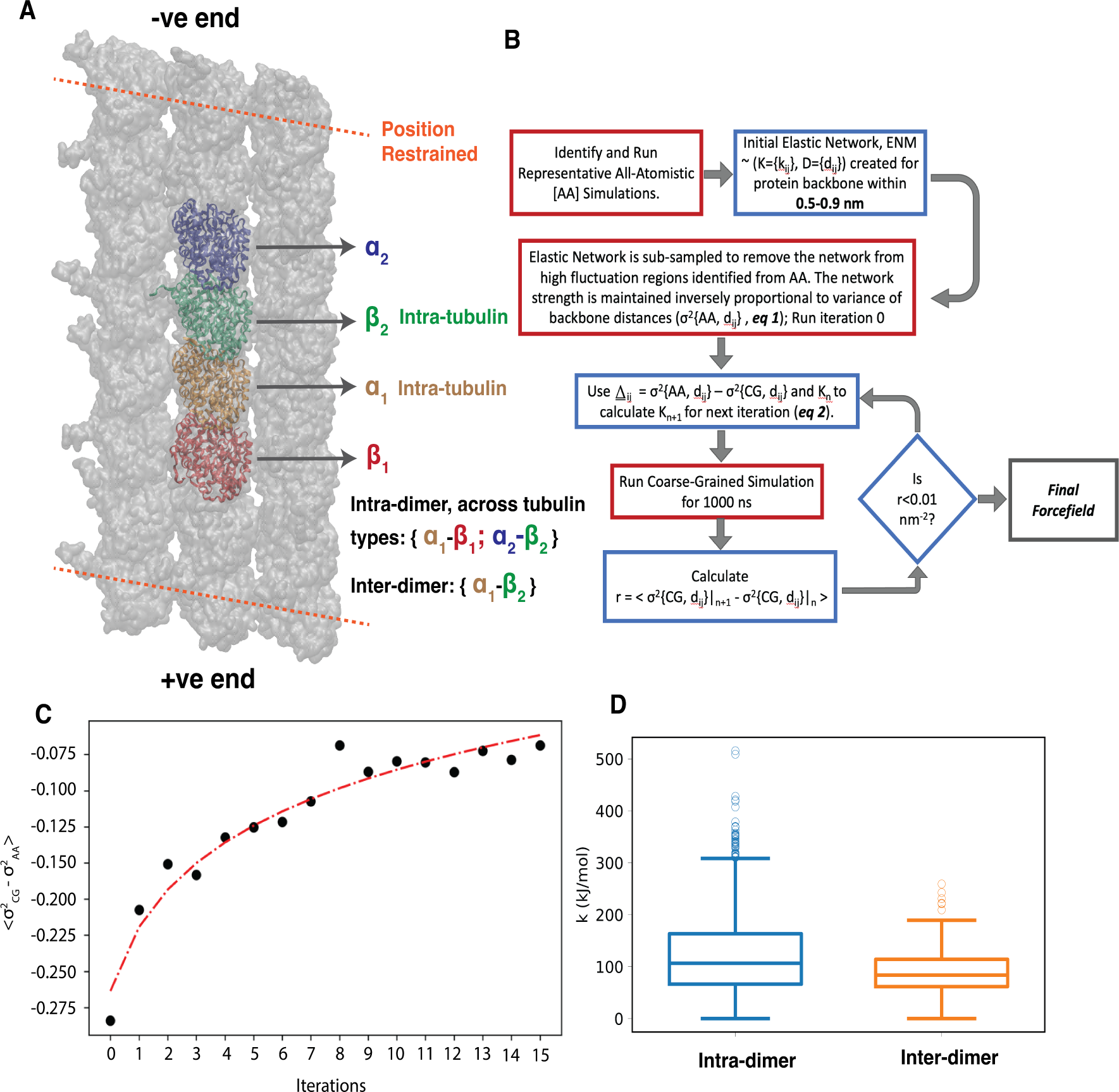
Parameterizing a more stable microtubule. *(A)* The all-atom microtubule patch used to parametrize the CG ENM. The tubulin monomers used for mapping fluctuations are highlighted. *(B)* Flowchart detailing steps for iterative optimization of elastic network. *(C)* Average deviation in fluctuation between coarse-grained and all-atom simulations over iterations. *(D)* Distribution of elastic network strength for inter-dimer and intra-dimer contacts.

### Inferring the elastic network with an iteratively-refined distance-based approach (IDEN)

The protocol (Iteratively-refined distance-based Elastic Netwrok — IDEN) encodes the dynamics of the microtubule patch into the elastic network in two steps — first, by removing highly dynamic regions from the ENM, and second, by iteratively calibrating the ENM through appropriately scaling the elastic network (Fig. 2A-D). Please refer to the Methods section for details of this parameterization. This optimization protocol reduced the number of harmonic restraints by 7.45% for inter-dimer contacts (average spring constant 119.8 kJ/mol); and 7.8% for intra-dimer contacts (average spring constant 94.6 kJ/mol). Fig. 1D shows the relative improvements of using the refined network compared to the out-of-box networks. This allowed for some local frustration to be relieved in this large assembled structure, leading to an overall more stable microtubule model.

### Bottom-up structural validations

In this section we present a set of local structural validations of our coarse-grained microtubule model by comparing local structural features from the full coarse-grained microtubule simula-tions with the atomistic simulations of the three-protofilament microtubule patch that was used for parametrizing our elastic network. Since, position restraints were applied to simulate a stable microtubule in atomistic simulations, we used the farthest tubulin dimer from the ends for our analysis to minimize the effects of these restraints (Fig. 2A). The last 100 ns of AA simulations were used for analysis, and was compared with last 500 ns of CG simulation, aggregated across three independent replicas. For the CG microtubule, only the central dimers from protofilament numbers 4 to 10 are used to minimize the effects that microtubule tips and seam might impart. Finally for the contacts at the microtubule seam, we used the central dimers of protofilament number 1 and 14, which constitute the microtubule seam.

Fig. 3A-C presents a comparison between local structures at AA and CG resolutions. As expected, the C-terminal tails of both *α*- and *β*-tubulins exhibit minimal high-probability distal contacts, which can be attributed to their disordered nature. Moreover due to high negative charges on individual tails, we did not observe any inter-tail contacts for both AA and CG simulations, with CG simulations exploring a larger set of contacts. Comparing between the two resolutions (Fig. 3A,B) reveal that discrepancies are mostly at the diagonal of the heatmap for both the tubulin types, referring to sequentially close contacts, which could be attributed to the compacting effects of the Martini forcefield. However, we do not observe any long-lasting interactions or inherent structure within the tails.

**Figure 3:**
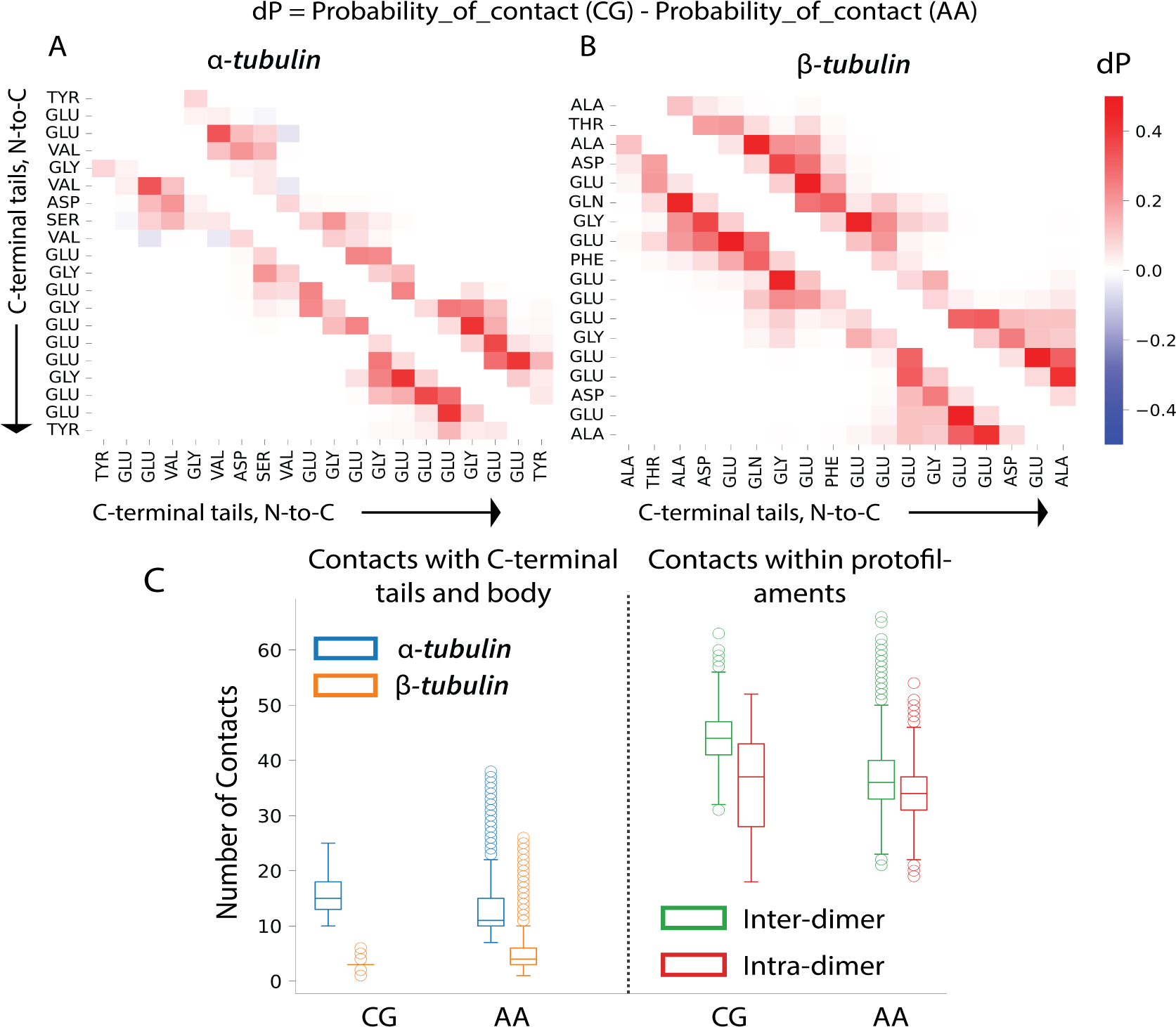
Bottom-up structural validations of our Martini microtubule. Heatmaps showing differ-ence is contact probabilities across AA and CG simulations for C-terminal tails in *α*-tubulins (*A*) and *β*-tubulins (*B*). (*C*) Comparisons of the number of contacts across AA and CG simulations. On the left are comparisons between the C-terminal tail and the body of the tubulin. On the right are comparisons between inter-dimer and intra-dimer contacts within a single protofilament.

Recent work have reported that interaction between C-terminal tails and the tubulin body can have both structural and functional relevance.^31^ Here we quantified the number of contacts between the tubulin tails and body (Fig. 3 C). Interestingly, the number of tail-body contacts is significantly (about three fold) higher for *α* tubulin compared to *β* tubulins even though the number of residues on C-terminus considered for this analysis was equivalent for both tubulins.

Along a single protofilament, the number of inter-dimer contacts were higher than intra-dimer contacts. This could be due to discrepancy between the nucleotide states at both the interfaces in GDP microtubules. These trends for the tail-body contact and tubulin dimer contacts were replicated in our coarse-grained simulations as well. Now, across protofilaments, while highly dynamic, we found an overall higher number of contacts for *α* tubulins compared to *β* tubulins (Fig. S1). Since the number of tubulin contacts across neighboring protofilaments were varied and depended on their relative position across protofilament number and distance from the ends, we have not reported the corresponding for our coarse-grained simulations. Finally, the cross protofilament contacts at the seam follows the same trend, with more *α*-tubulin contacts compared to *β*-tubulin (Fig. S2).

### Bio-mechanical validations — axial bending modulus

Microtubules are among the primary structural elements of the cell and their mechanical prop-erties have been extensively investigated — both in isolation and in presence of physiolog-ical/simulated stimulus. ^11,12,36^ For example, bending modulus has been measured either by directly pulling on the microtubules with optical tweezers or through fitting equilibrium ther-mal fluctuations to mathematical models (both linear and non-linear). ^12,37^ These studies have resulted in a range of reported values spanning about an order of magnitude depending on nucleotide and/or other ligand associated states. But the consensus observation is that the microtubules are highly rigid biological assemblies that are capable of withstanding extensive mechanical stress and remodelling other cellular assemblies like membranes.

Therefore, in order to extend our structural models to more complex physiological settings we need to be able to reproduce these elastic features. To measure the axial Young’s modulus (*E_a_*) of our coarse-grained model microtubules, we created a periodic microtubule (∼10 tubulin dimers long) along the microtubule axis and simulated it with applied anisotropic pressure along the axis. (*P_z_*, Fig. 4 A). The slope of straight-line fit, assuming an elastic regime, in our inferred strain-strain plot resulted in *E_a_* = 0.64 ± 0.08 GPa (Fig. 4 B). Further, flexural ridgidity (*γ* = *E_a_I*) can be inferred using the second moment of cross sectional area (I). Our simulations suggest a flexural ridgidity of 12.6*pNµm*^2^. These reported values are close to experimental reports and within the reported ranges (*E_a_* ∼ 0.6 − 1.1GPa and *γ* ∼ 1.36 − 35.8*pNµm*^2^; Fig. S2).^12,37^

**Figure 4:**
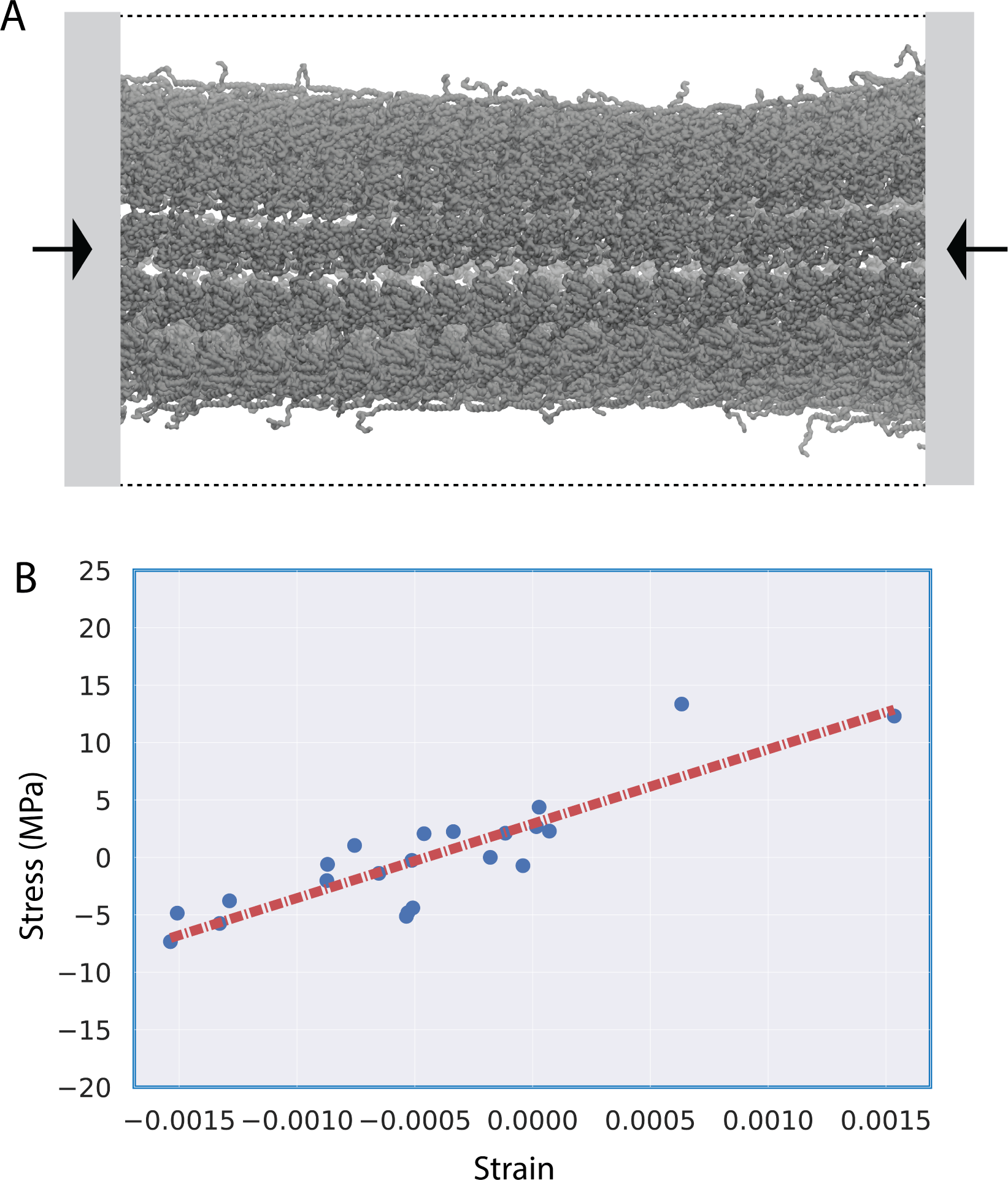
Simulating a microtubule that interacts with its periodic image establishes this micro-tubule exhibits mechanically correct properties. *(A)* A representative snapshot of a microtubule under applied axial pressure (1 bar). *(B)* Stress-Strain curve calculated from applied pressure simulations. The slope of the regression line is the axial Young’s modulus (*E_a_*).

### Structural Analysis of GDP-Bound Microtubules with Optimized ENM

In this section, we highlight some preliminary structural observations from our simulations of GDP-bound microtubules (∼ 200 nm long, 25 tubulin-dimers, ∼ 6 million interaction-centers; Fig 5 A). For these analyses, we have aggregated the trajectory data from three independent replicas, each run for 1 *µs*. The simulations successfully replicated local bending phenomena, at the extremities of microtubules — driven by thermal fluctuations, whilst preserving the microtubule’s intrinsic cylindrical form (Fig. 5 A-C).

**Figure 5:**
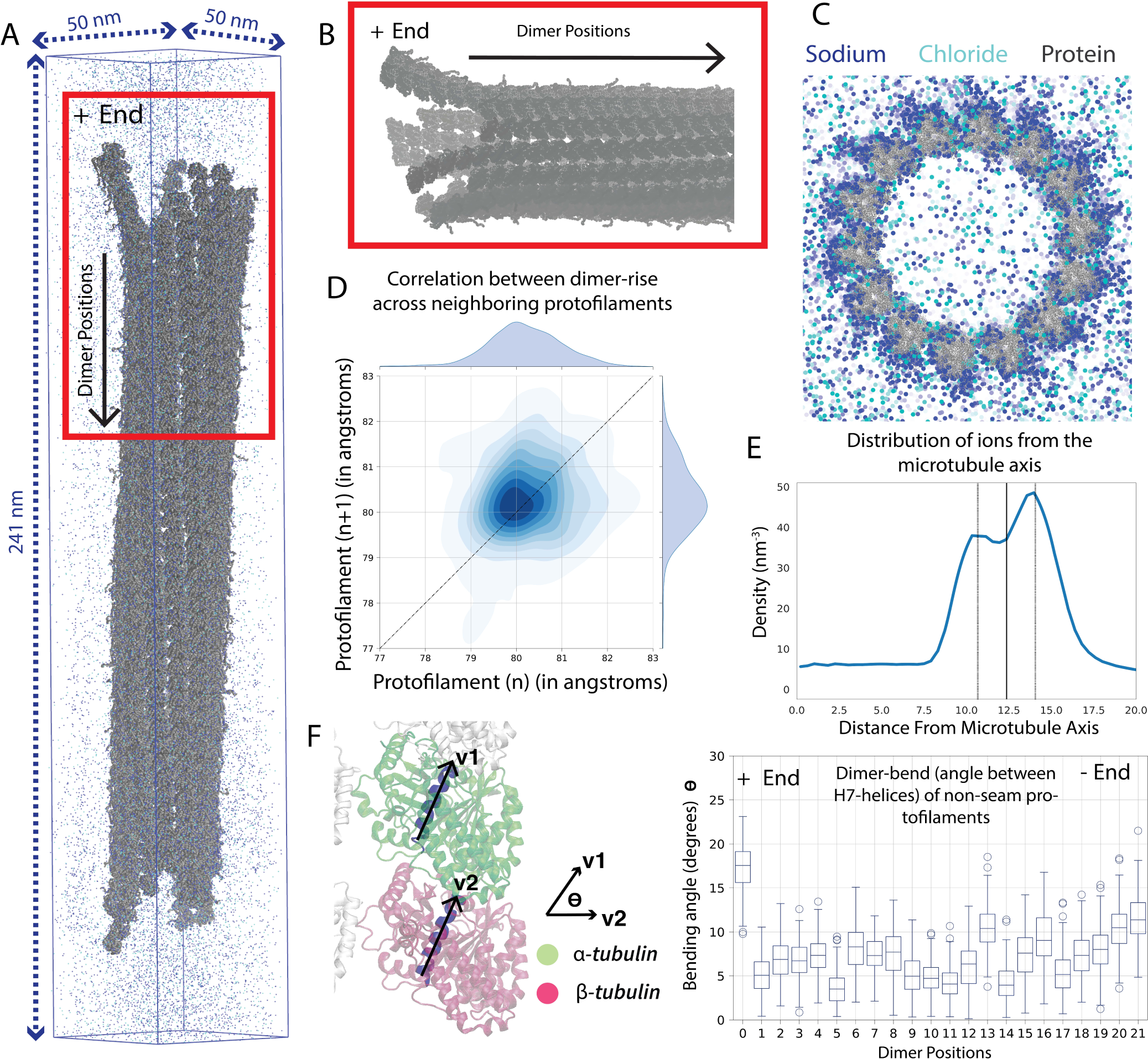
Structural analysis of 200nm microtubule simulations. A -Representative snapshot of the whole microtubule, with ions shown for reference. B -Representative snapshot of the microtubule plus end. C -Axial view of the microtubule with ions. D -Contour plot showing correlation between the inter-dimer distances of adjacent protofilaments. E -Density distribution of sodium ions from the microtubule axis. F -Angle made by regression lines representing H7-helix of *α* and *β* tubulin over microtubule length for a central protofilament

Similar to the observations by Igaev *et al.*, our study also identified a modest correlation (pearson’s r ∼ 0.34) in the spacing between adjacent dimers (*α*-*β*) across neighboring protofilaments of the microtubule. ^18^ However this effect was varied across protofilament number and time, depending on whether it forms contacts across protofilaments (Fig. 5 D). This microscale interaction could be pivotal for influencing alterations in the lattice configuration and modulating the localized interplay between the microtubule and its binding proteins, including motor proteins, which depend on the spacing within the lattice structure for their functions.^38^Moreover, this could also speak to the relative stabilities of different nucleotide associated states or tubulin isoforms and how perturbations in lattice travel along the microtubule.

Given the explicit-solvent nature of our simulations, we can also report local enrichment of solvated ions. Fig. 5E is the normalized probablility of distances of ions measured from the micro-tubule axis. We observed an increased partitioning of monovalent ions to densities greater than 5 times the bulk-density at the microtubule surface. Particularly, the density of ions partitioned at the outer-side was higher than the inner. This could be due to highly charged disordered C-terminal tails on the tubulins that contribute to local accumulation of ions. Moreover, we observe an ion screening distance (debye-length) of close to 1.5 nm, similar to previous estimates from experiment. ^39^

Finally, we calculated the bending/curving of individual dimer through the angle made by the H7-helix of *α* and *β* tubulins (Fig 5F). H7-helix is a conserved core helix of tubulins that has contacts with the nucleotide-binding domains, and has been previously used to characterize dimer bending.^40^ Fig. 5F, shows the distribution of the angles of a central protofilament (combined data from protofilament 7 and 8) and the microtubule seam (combined data from protofilament at the seam — 1 and 14) respectively. In both cases, an enhanced bending at the microtubule termini was observed, which is likely attributed to the reduced stabilizing interactions between adjacent protofilaments at these ends. Please note here that interactions between adjoining protofilaments can contribute to multiple protofilaments peeling off together, as noted from prior all-atom simulations of microtubule tips. ^19^ These preliminary observations suggest a stable microtubule lattice that reproduces local structural and dynamic patterns previously reported in literature.

## Discussion and Conclusions

In this work, we introduce a Martini 3.0 microtubule capable of stably producing a 2*µs* simulation of a ∼200nm GDP-bound microtubule, while maintaining the generalizability of the Martini forcefield. This choice provides the ability to not only easily add other proteins to the simulation box, but includes the residue level detail required for simulation of explicit solvent around the microtubule and the highly charged disordered C-terminal tails.

Here, the elastic network is iteratively parametrized to reproduce distance fluctuations in atomistic simulations of microtubule patches. Our methodology employs a bottom-up coarse-graining strategy originally introduced by Globisch *et al.* ^35^ and maintains fidelity to the original Martini Hamiltonian while introducing modifications to the force field via the elastic network, which is effectively a correction term. Therefore, this model leverages all the benefits afforded by the Martini forcefield such as a amino-acid sidechain level specificity, large library of parametrized interaction partners (proteins, small molecules, lipids, and metabolites), high transferability, and easy optimization/modification.

The Martini forcefield has been extensively adopted by the research community to accurately study macromolecular systems, with diverse components at large spatio-temporal scales. For example, the forcefield has been employed to construct intact severe acute respiratory syndrome coronavirus 2 (SARS-CoV-2) virion, with an accurate representation of membrane, trans-membrane proteins and N-bound RNA fragments. ^41,42^ Recently Mosalaganti et al. probed the structure and dynamics of the nuclear pore complex (∼ 120*MDa*) which features multiple copies of thousands of distinct proteins forming a torriodal scaffold. ^43^ Similarly, Javanainen *et al.*, investigated the rotational and translational diffusion of proteins in crowded membranous environments. ^44^ A common feature in these large systems is how micro-scale, local interactions compound, resulting in macro-scale and/or mechanical changes — bridging information across multiple scales. Therefore, supplementing this popular knowledge-based forcefield with a bottom-up coarse-graining approach that encodes local micro-scale interactions while preserving macroscale features, addresses an important challenge in multiscale modelling.

Specifically in the context of microtubules, along the similar vein as this work, Martini enhanced with IDEN also makes it possible to study other nucleotide associated states (GTP and GMPCPP) and tubulin isoforms/constructs at larger spatiotemporal scales.^45–47^ This approach is particularly helpful in alleviating some key drawbacks of the MARTINI model regarding accuracies of nucleotide and/or small-molecule forcefield by implicitly encoding their effect in form of local fluctuations.

Beyond microtubules, this also enables understanding the diffusion and binding of several microtubule-associated proteins (such as Tau, PRC1 and EB1) and other small molecules like taxol on microtubule lattice. ^48–54^ For example, tau — an important microtubule binding protein associated with the Alzheimer’s disease, has been recently shown to form condensates while interacting with the microtubule, thereby rearranging local dimer separation, which can in turn rearrange neighboring protofilament architecture, as evidenced from correlated dimer-separations on adjoining protofilaments in Fig 5D. ^48^

Another potential area for future research could involve assessing the influence of changing salt concentrations on microtubule dynamics. In our research, we observed localized accumulation of sodium ions in the outer regions of the microtubule, especially near the tubulin C-terminal tails. Previous research has documented the effects of varying salt concentrations on microtubule bundling and interactions between microtubules. Employing an explicit-solvent model capable of capturing local salt concentration fluctuations, this coarse-grained microtubule model could significantly contribute to advancing our understanding in those areas.

Finally, such modelling approaches can be helpful for the experimental structural biology community. For example, our simulations can serve as templates to identify micro-structures in cryo-electron tomographs. This can be particularly useful for characterizing defects in microtubule lattice and highly flexible microtubule tips.

Since this model builds on the massively popular Martini model to derive its benefits, it also inherits Martini’s problems. Even with significant reparameterizations in the newest iteration of the forcefield, there are still reported problems with protein-protein interactions. This is particularly relevant for disordered regions. As an important functional region of the microtubule is disordered — C-terminal tails, this inability to capture accurate conformational space is an important limitation. While, in this work we have been particularly careful in reproducing this space to the accuracies comparable to AA simulations through re-scaling of protein-water interactions, Martini 3 still generated a somewhat compacted conformation evidenced by increased self-interactions within each tail. Moreover, even with a significantly reduced elastic network model to allow for more fluctuations, this model cannot still capture large conformational transitions such as folding/unfolding of domains. Another important limitation of the forcefield is the inaccurate entropy-enthalpy balance. Considering the optimization strategy employed for Martini forcefields involves balancing interactions between coarse-grained interaction centers to reproduce accurate free energies, this can lead to inaccurate entropy-enthalpy balance. Therefore, we need to be careful in considering kinetic information, and primarily focus on comparisons rather than absolute kinetics.

Finally, with classical molecular dynamics forcefield, it is not possible to capture active processes that are often associated with the microtubules such as GTP hydrolysis and walking of motors without careful additional modifications. Beyond the forcefield, another challenge in this approach was appropriately modelling the microtubule seam. Here. we inferred the structure by rotations and translations from the cryo-EM lattice. Therefore, we do not accommodate for any conformational variances or specific geometry at the seam, considering unavailability of structural data.

Future work in our group aims to address some of the limitations — particularly simple structural transitions and some reactive processes with alternatives of elastic-network that allow spontaneous forming and breaking of bonds.^55–58^ A recently developed proxy for elastic net-work are *GO̅*-like models that encode conformational dynamics more effectively, similar to *GO̅*-Martini. ^59,60^ This will possibly help in capturing more disordered states at microtubule tips, and binding-unbinding dynamics of tubulins. Furthermore, this will build a stronger bridge with thermodynamics through carefully parametrizing the *GO̅* model to reproduce experimental binding affinities.

To summarize, this work presents a first Martini microtubule, with all the benefits and drawbacks of that choice of forcefield, as well as the scheme used to make this microtubule stable. We believe this will provide a valuable stepping stone for other CG-MD simulations seeking to characterize the molecular biophysics dictating microtubule function, including but not limited to solvent effects, protein binding partners, and microtubule lattice dynamics. Finally, this approach to microtubule molecular dynamics can also be used in tandem with cryo-electron microscopy to understand the intrinsic dynamics of microtubules and their binding partners seen in various relevant and well-studied *in vitro* and *in vivo* environments. ^61,62^

## Materials and Methods

### CG Simulation Protocol

Sodium chloride at 150 mM concentration was introduced to solvated protein systems, followed by energy minimization via steepest descent until achieving machine precision. Subsequently, an initial equilibration phase of 100 ns (timestep: 2 fs) was undertaken in the isothermal ensemble, with protein backbone beads restrained (force constants: 1000 kJ/mol). Three additional equilibration steps for 100 ns each followed, incrementally increasing timesteps (2 fs, 5 fs, 10 fs) in the isothermal-isobaric ensemble, while keeping the protein backbone restrained. Throughout equilibration, a v-rescale thermostat and Berendsen barostat maintained constant temperature (300 K) and pressure (1 bar). Ultimately, a one micro-second production simulation with a 10 fs timestep, Parinello-Rahman barostat, coupling time constant of 12 fs, and compressibility of 3.0 × 10*^−^*^4^/bar was run. ^63^ For all the equilibration, and production steps, the electrostatic cutoff distance is set at 1.6 nm, and to account for long-range electrostatic interactions, we employ the particle-mesh Ewald (PME) method. ^64,65^

### All-Atom Simulation Protocol

All the all-atom simulations were performed with gromacs 2022.4 with CHARMM36m/TIP3P forcefield.^66,67^ The initial model of a microtubule patch was downloaded from the cryo-EM structures deposited in the protein data bank —-6DPV for GDP microtubule. ^25^ The microtubule patches are elongated along the microtubule axis, to twice the length of deposited structures. The C-terminal tails and other smaller internal missing loops were added to the structure with CHARMM-GUI.^68^ The initial structure was passed through the CHARMM-GUI solution builder to generate the input files for the simulation. Water and 150 mM sodium chloride is added to the simulation system, before energy minimization with stochastic gradient descent to reach machine precision. The system was equilibrated for 1 ns with isothermal, isochoric ensemble using v-rescale thermostat at a coupling time of 1 ps. Following this, the system was further equilibrated in a isothermal, isobaric ensemble for 5 ns. For all the equilibration steps, the protein backbone is restrained allowing for side-chain and solvent to equilibrate.

In our production simulations, strategic position restraints were applied to distinct microtubule segments, allowing us to focus on specific regions of interest. Within the microtubule lattice, we imposed position restraints on the backbone of the two terminal tubulin monomers, with a force constant of 10 kJ/mol. Meanwhile, solely the tubulins located at the minus end of the microtubule were subject to position restraints for microtubule ends. The tubulin contacts situated farthest from the restrained tubulins were employed to derive the corresponding elastic network.

### Iteratively Refined Distance-based Elastic Network (IDEN)

Our approach for refining ENM involved first defining a list of contacts *D* = *d_ij_*, the distance between non-neighboring C-*α* atoms in the PDB structure, was within 5-9 Å (*L_cutoff_*). This list was further sub-sampled by removing outliers, where the experimental values from the PDB-derieved structures are outside three standard deviation of the corresponding probability distribution — derieved from the atomistic simulations of the patch, assuming a local normal distribution. This sub-sampling explicitly removes the ENM from highly dynamic parts of this biomolecular assembly.

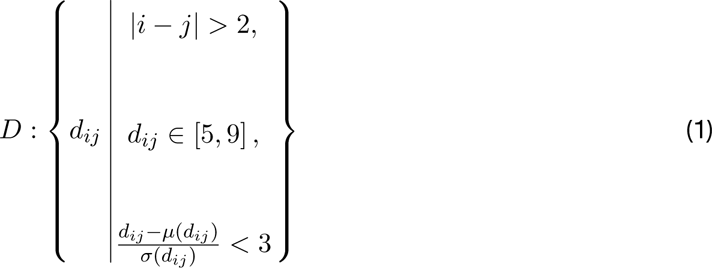

The elastic network strength (*k_ij_*) is initially constructed inversely proportional to the variance of respective distances (*d_ij_*).

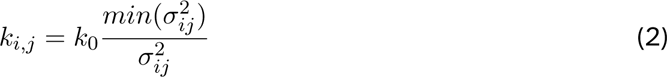

where, *σ*^2^ refers to the variance of contact distances; and *k*_0_ = 500 kJ/mol.

The initial atomistic model is then coarse-grained using the Martini 3 mapping scheme. Following minimization and equilibration, the production simulation is run for 1 micro-second with position restraints on the end tubulins to mimic the atomistic simulations. The generated trajectory can now be processed to infer the elastic-network part of the forcefield. We have followed the iterative optimization stratergy originally introduced by Globisch *et al.* to match the variance of coarse-grained distances to the all-atom simulations with the following update rule.

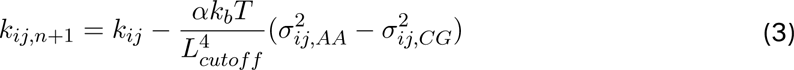

where, n refers to the iteration number, 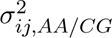 is the variance in contact-distances, *α* is a constant and in this work we have used 1050, similar to Globisch *et al.* ^35^ For the iterations, we tracked 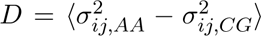 over the last 500 ns of simulation time. The optimization was stopped when the update led to minimal/no improvement in D (Fig. 2 C).

### Microtubules under Axial Pressure

To compute the flexural rigidity of our model microtubules, we followed closely the protocol outlined by Igaev *et al.* ^18^ First, we created a smaller microtubule (∼ 90 nm), and modified the box such that the microtubule is periodic along the axis (Z). The system was solvated and 150 mM ions were added. The initial equilibration followed the exact same protocol as for our other coarse-grained simulation, but with anisotropic pressure coupling (*P_x_* = *P_y_* ≠ = *P_z_*). Finally, an additional long equilibration simulation was performed to equilibrate the simulation box along the microtubule axis, due to the additional axial force. The applied axial pressure, *P_z_* ranged from −0.25 bar to +2 bar. With values greater than +2 bar, the cylindrical architecture of the microtubule started falling apart. And at values below 0.25 bar, the microtubule was not able to maintain periodic contact, due to the elastic network. We acknowledge that going away from an elastic network model towards a *GO̅*-like model here could be more accurate because that would allow us to then simulate microtubules at much higher values of extension and compression.

Therefore the net stress, *σ_zz_* can be computed as —

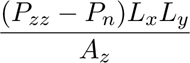

where, *L_x_* and *L_y_* are box dimensions; *P_z_z*, *P_n_* is the measured axial and normal pressure, and *A_z_* is the cross-sectional area of the microtubule. We calculated 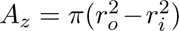 and the second moment of cross-sectional area 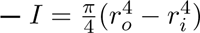, by first inferring the inner (*r_i_*), and outer (*r_o_*) radius of the microtubule using distribution of protein-backbone distances measured from the microtubule axis.

The strain is computed as,

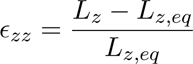

where *L_z,eq_*is the mean axial box length.

The last two micro-second of simulation-time was used for inference, with block averaging through 200 ns windows for error analysis. This analysis used a net sampling time of ∼ 8*µs*

## Acknowledgements

We would like to thank a number of people who have been helpful at various stages of this work, and continue to help influence future directions: Reza Farhadifar, Pilar Cossio, Dan Needleman, Will Conway, Michael O’Brien, Christopher Edelmaier and Adam Lamson. The Flatiron Institute is a division of the Simons Foundation.

## Data and Code Availability

The scripts for IDEN adapted for microtubules are shared here https://github.com/flatironinstitute/martini-microtubule. The complete trajectory will be shared shortly in a publicly available repository.

## Supplementary Information

### Supplementary Figures

**Supplementary Figure 1:**
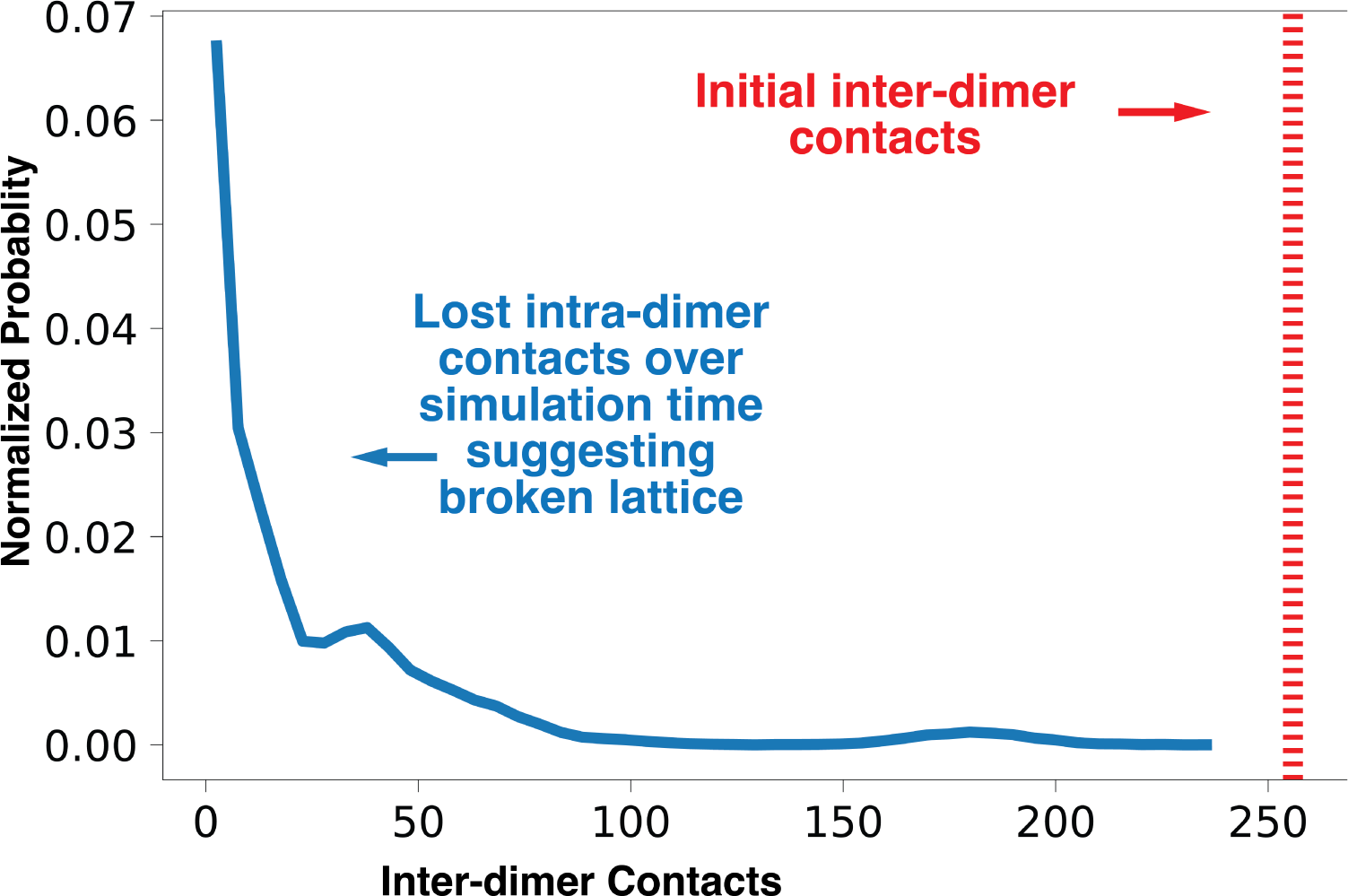
The probability of inter-dimer contacts after 400 ns of simulation time, with no inter-dimer elastic network.

**Supplementary Figure 2:**
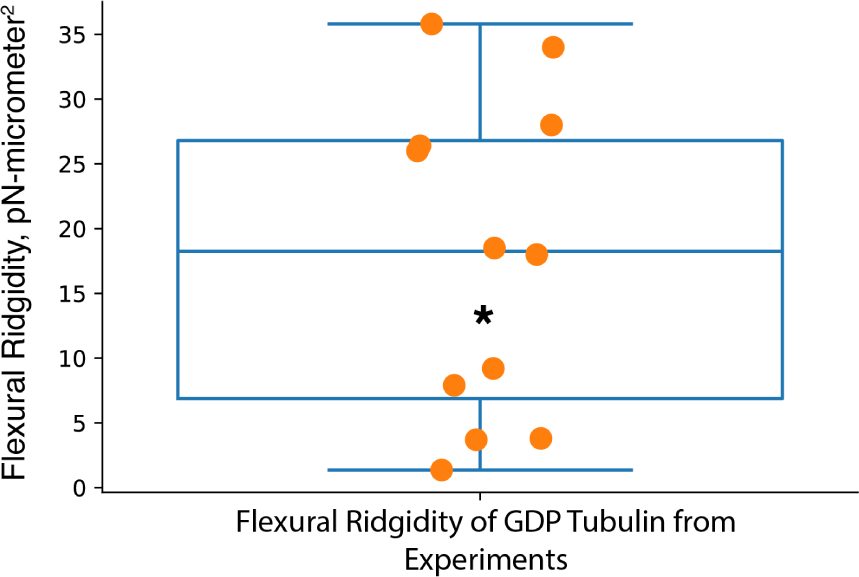
Flexural ridgidity of GDP tubulins reported from experiments. ‘*’ marks the flexural ridgidity reported in this work. Data adapted from ^12^.

**Supplementary Figure 3:**
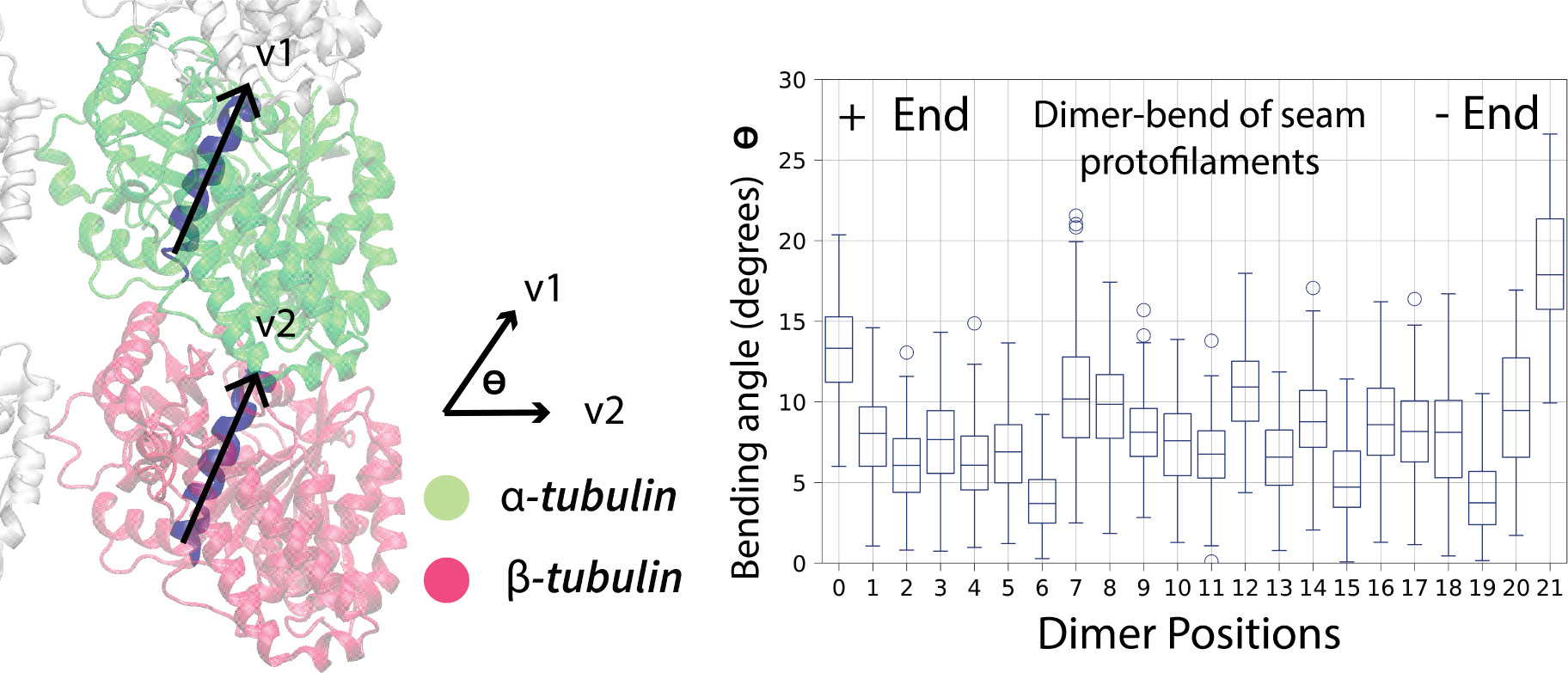
Tubulin bending of the protofilament at the **seam**, characterized by the angle that H7-helix of *α*, and *β* tubulins form.

### Supplementary Video

1 micro-second trajectory of a full microtubule.

